# Simian Immunodeficiency Virus and Storage Buffer: Field-friendly preservation methods for RNA viral detection in primate feces

**DOI:** 10.1101/2023.08.28.555131

**Authors:** Tessa H.C. Wilde, Rajni Kant Shukla, Christopher Madden, Yael Vodovotz, Amit Sharma, W. Scott Mcgraw, Vanessa L. Hale

## Abstract

Wild non-human primates carry many types of RNA viruses, including simian immunodeficiency virus (SIV), simian foamy virus, simian T-cell leukemia virus, and hepatitis C virus. These viruses can also infect humans via zoonotic transmission through handling and consumption of primate bushmeat. Characterizing viral prevalence and shedding in natural hosts is critical to understand infection and transmission risks within and between primate species. Here, we sought to identify a robust “field-friendly” method (i.e., without freezing or refrigeration) for preserving viral RNA, specifically SIV, in primate fecal samples. Fecal samples were collected from a mantled guereza colobus (*Colobus guereza*) housed at the Columbus Zoo and Aquarium. Samples were homogenized and inoculated with three concentrations (low, medium, high) of inactivated SIV virus and preserved in four different storage buffers (DNA/RNA Shield, RNA*later*, 95% Ethanol, and Viral Transport Medium). SIV viral RNA was then extracted from samples at four time points (1 week, 4 weeks, 8 weeks, and 12 weeks) to determine the efficacy of each buffer for preserving SIV RNA. Quantitative RT-PCR was used for detection and quantification of viral RNA. At all concentrations, DNA/RNA Shield yielded the highest average SIV virion concentrations. We then successfully validated this approach using fecal samples from known SIV-positive and SIV-negative sooty mangabeys (*Cercocebus atys*) housed at Emory National Primate Research Center. Our results indicate that DNA/RNA shield is an optimal “field-friendly” buffer for preserving SIV RNA in fecal samples over time, and may also be effective for preserving other RNA viruses in feces.

**Importance:** Human immunodeficiency virus (HIV) was introduced into human populations through zoonotic transmission of SIV from African primates, leading to a global epidemic and ongoing worldwide public health issue. SIV occurs naturally in over 40 primate species in sub-Saharan Africa and these viruses have crossed species barriers on multiple occasions, leading to the spread of HIV-1 and HIV-2. Quantifying RNA viruses in wild primate populations can be challenging as invasive sampling is often not feasible, and many field stations lack ready access to a freezer for storing biological samples. This study compares SIV RNA preservation and recovery across multiple storage buffers to identify a robust field-friendly option for RNA viral detection in noninvasively collected feces. Our results will inform future fieldwork and facilitate improved approaches to characterizing prevalence, shedding, and transmission of RNA viruses like SIV in natural hosts including wild-living nonhuman primates.

## Introduction

Wild non-human primates can carry many types of RNA viruses, including simian immunodeficiency virus (SIV), simian foamy virus (SFV), simian T-cell leukemia virus (STLV), and hepatitis C virus (HCV). These viruses can also infect humans via zoonotic transmission through handling and consumption of primate bushmeat. Characterizing viral prevalence and shedding in natural hosts is critical to understand infection and transmission risks within and between primate species. Human immunodeficiency virus (HIV) was introduced into human populations through zoonotic transmission of the SIV from African primates, leading to a global epidemic and ongoing worldwide public health issue (Hahn et al., 2000). SIV occurs naturally in over 40 primate species in sub-Saharan Africa and these viruses have crossed species barriers on multiple occasions, leading to the spread of HIV-1 and HIV-2 in humans populations (Apetrei et al., 2004). Humans continue to be exposed to these RNA viruses through handling and consumption of primate bushmeat, so it is important to investigate wild-living nonhuman primate populations to understand SIV prevalence, shedding, and transmission in natural hosts (Aghokeng et al., 2010; Ahuka-Mundeke et al., 2017). Importantly, the COVI9-19 pandemic also made us globally aware of the value and need for RNA virus surveillance in wild animal populations to assess health risks to humans and to animals (Delahay et al., 2021; Ehrlich et al., 2023; Gwenzi et al., 2022; Hale et al., 2022).

Assessing presence and abundance of RNA viruses in wild primate populations can be challenging as invasive sampling is often not feasible, and many field stations lack ready access to a freezer for storing biological samples. Ling and colleagues (2003) were one of the first groups to investigate the sensitivity of SIV detection in primate fecal samples. They tested fecal samples from laboratory-housed sooty mangabeys (*Cercocebus atys*) for SIV viral RNA and determined that RT-PCR detected the presence of SIV viral RNA in fecal samples from about 50% of the positive mangabeys. They went on to test this method in the field and were able to confirm one case of SIV, out of 61 samples, in a population of wild-living sooty mangabeys in Sierra Leone (Ling et al., 2004). Their samples were frozen in the field, a process not possible in every field situation.

Additional studies have focused on red colobus monkeys (*Piliocolobus badius*) and sooty mangabeys (*Cercocebus atys*) in Taï National Park, the likely site of origin for the HIV-2 epidemic (Locatelli et al., 2011; Santiago et al., 2005). By assuming that RT-PCR can detect SIV in about 50% of fecal samples, these authors estimated between a 50-60% prevalence of SIV was predicted in wild populations of these two primate species. Both studies tested only for the presence or absence of the SIV virus and utilized *RNAlater* (Thermo Fisher), a storage buffer that allows fecal samples to be stored at ambient temperature for a short period of time.

*RNAlater* is increasingly used to store fecal samples in the field; however, according to *RNAlater*’s guidelines, RNA quality may begin to degrade after just one week of storage, if not frozen. Many field studies involve weeks or months in the field, so testing storage methods with a range of ambient stability times is useful for researchers lacking freezer access in the field.

Here, we tested multiple “field-friendly” methods (i.e., those not requiring freezing or refrigeration) for preserving viral RNA, specifically SIV, in primate fecal samples. The storage buffers we tested included: DNA/RNA Shield, RNA*later*, 95% Ethanol, and Viral Transport Media (VTM). Fecal samples in each buffer were inoculated with three concentrations of SIV virus: 300,000 virions/mL (high), 30,000 virions/mL (medium), and 3,000 virions/mL (low). These concentrations fall within the range of SIV concentrations detected in the plasma of naturally-infected SIV-positive primates (<500 copies/mL to greater than 2×10^6^ copies/mL (Ling et al., 2003)). SIV viral RNA was then extracted from samples at four time points (1 week, 4 weeks, 8 weeks, and 12 weeks) (**Figure 1**). We aimed to determine which buffer was the most effective at preserving viral RNA at different time intervals. Additionally, we characterized (a) the threshold of detection and (b) quantified the amount of SIV viral RNA present in our study samples.

**Figure 1:**
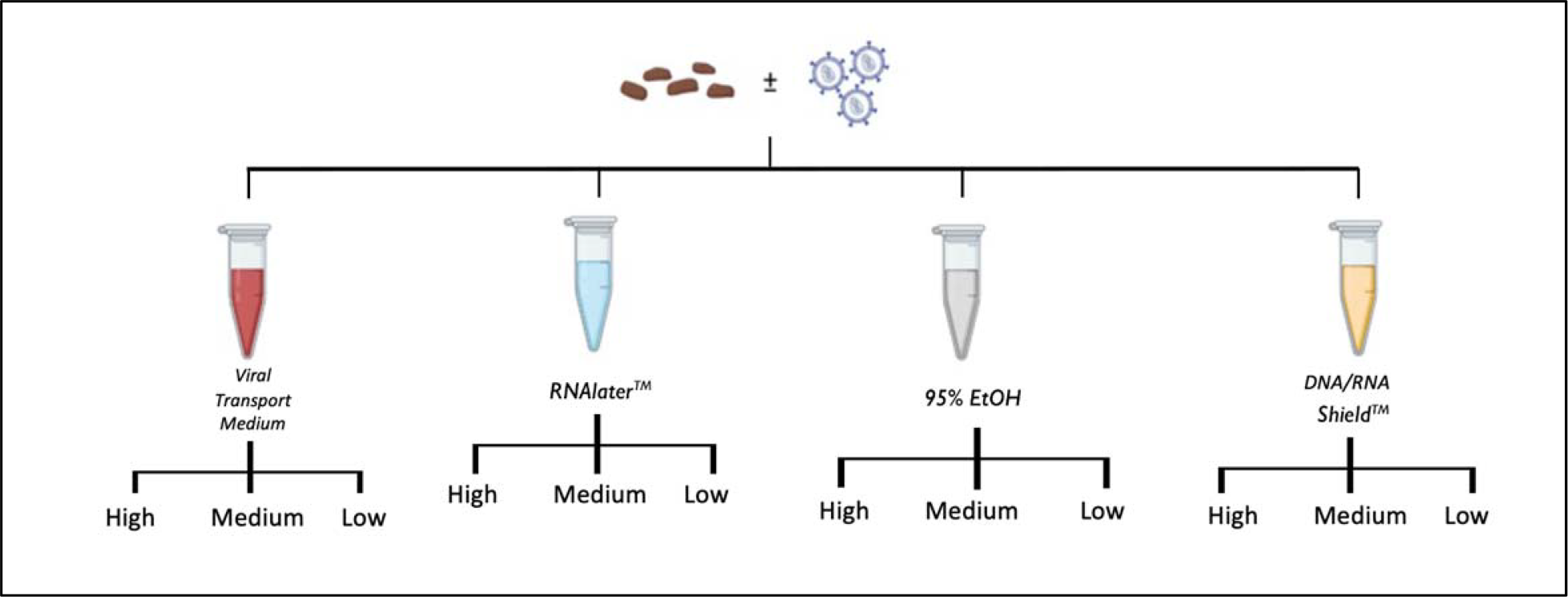
Experimental design. A single homogenized fecal sample from a colobus monkey was aliquoted into four different storage buffers (Viral Transport Medium, RNA*later*, 95% Ethanol, and DNA/RNA Shield) and inoculated with high (300,000 virions/mL), medium (30,000 virions/mL), or low (3,000 virions/mL) concentrations of SIV. RNA was then extracted from each sample at weeks 1, 4, 8, and 12.

## Results

### SIV virion concentrations by buffer

SIV virion concentrations varied significantly by storage buffer but not significantly over time (Kruskal-Wallis: buffer *p*=0.025; time *p*=0.322). These results were driven by the highest and lowest performing buffers which were DNA/RNA Shield and ethanol respectively (Dunn’s test: DNA/RNA Shield v. Ethanol: *p*=0.004; Ethanol v. RNA*later*: *p*=0.022; All other pairwise comparisons between buffers were not significant (**Table 1**)). At all SIV concentrations (high, medium, and low), DNA/RNA Shield preserved the greatest percentage of SIV virions, followed by RNA*later*, viral transport media, and ethanol respectively (**Figure 2**). For example, at the high SIV concentration, DNA/RNA Shield preserved 84% of the SIV virions at 12 weeks, while RNA*later* preserved only 45% of the SIV virions at 12 weeks.

**Table 1:** P-values from Dunn’s pairwise test comparing SIV virion percent yields by storage buffer. All concentrations and time points were combined in this analysis. Percent yields were based on the original amount of SIV spiked into each sample. *Significant p-value.

|  | **RNAlater** | **VTM** | **95% Ethanol** |
| --- | --- | --- | --- |
| **DNA/RNA Shield** | 0.556 | 0.148 | <0.004* |
| **RNAlater** | - | 0.409 | 0.022* |
| **VTM** | - | - | 0.124 |

**Figure 2:**
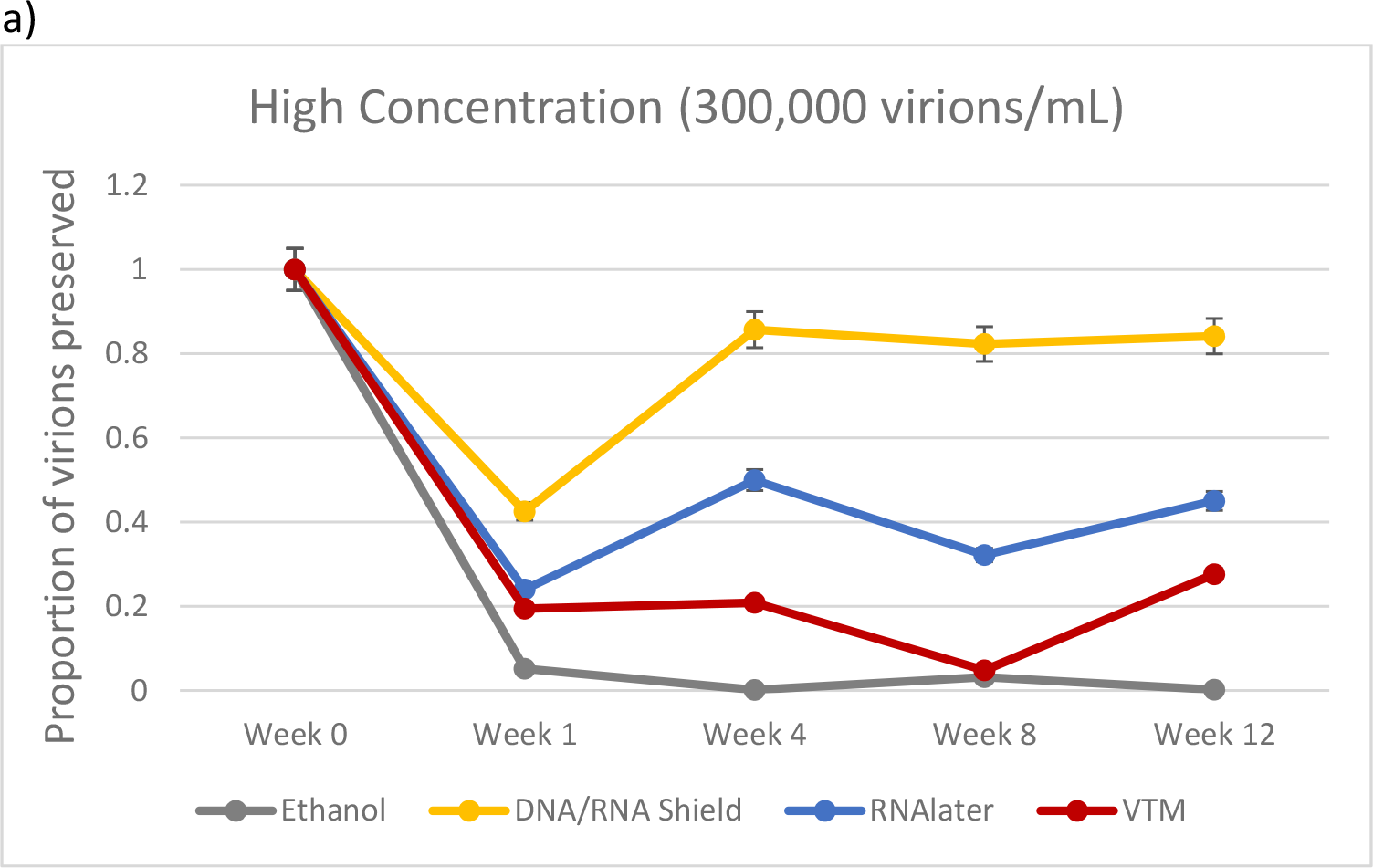
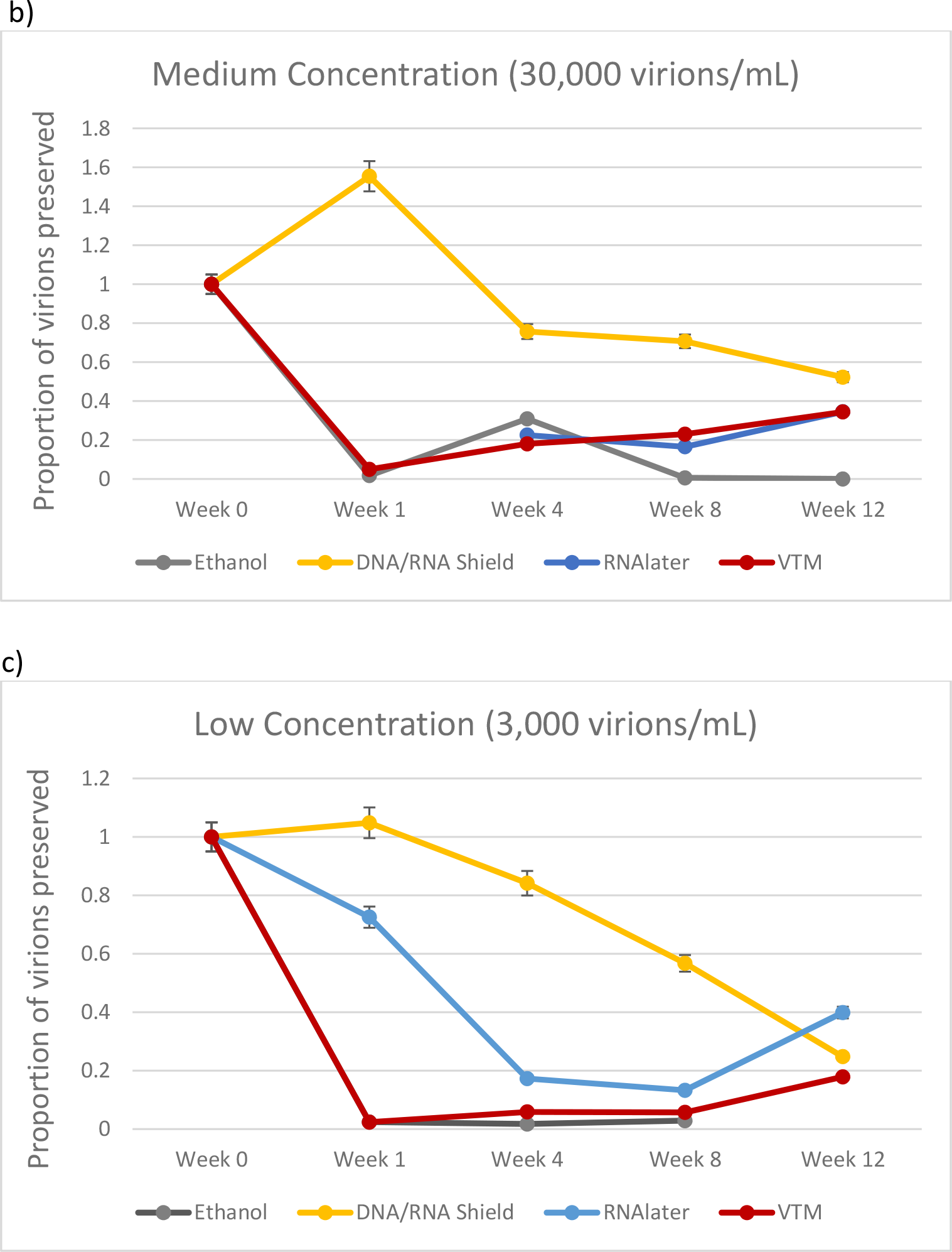
SIV yield by buffer and time. Percent yield of SIV virions (compared to spiked in viral load) extracted from fecal samples and quantified via RT-PCR. Fecal samples were spiked with (a) High (300,000 virions/mL), (b) Medium (30,000 virions/mL), and (c) Low (3,000 virions/mL) concentrations of SIV and preserved in four storage buffers (95% Ethanol, DNA/RNA Shield, RNA*later*, and Viral Transport Medium) for up to 12 weeks.

### SIV virion concentrations over time

We did not observe any significant differences in virion concentrations over time (Weeks 1-12) when we compared samples within and between buffers. However, our sample size was small, and we had limited power to detect potential differences. Notably, we did observe a non-significant decrease in virion concentrations over time in DNA/RNA Shield at medium and low, but not high, SIV concentrations. Importantly, we kept all samples (except VTM) at room temperature for 12 weeks. DNA/RNA Shield manufacturer guidelines indicate that samples can be stored at room temperature for up to 4 weeks, but should then be frozen at -80°C to prevent further sample degradation. SIV virion concentrations were relatively consistent over time across all other buffers. In a few Week 1 samples, we observed lower SIV virion concentrations as compared to subsequent timepoints (Weeks 4, 8, or 12).

### Validation cohort

To validate our approach, we then collected blood and fecal samples from SIV-positive (n=5) and SIV-negative (n=5) sooty managabeys housed at Emory National Primate Research Center (NPRC). Blood testing confirmed the SIV status of these 10 mangabeys. No SIV viral RNA was detected in the fecal samples of the SIV-negative mangabeys. In the SIV-positive mangabeys, we detected SIV viral RNA in two of the five monkeys (**Table 2**). The SIV viral load detected in both positive fecal samples was low (<1,000 copies/mg), and both fecal samples were associated with high serum viral loads (>120,000 copies/mL). SIV-positive managabeys in which no SIV viral RNA detected in feces all had low serum viral loads (< 32,000 copies/mL).

**Table 2:** Average Ct value and estimated SIV viral load for fecal and blood samples collected from SIV+ and SIV-sooty mangabeys (*Cercocebus atys*) at Emory National Primate Research Center.

| **Individual** | **SIV Status** | **Average Ct Value** | **Estimated fecal viral load (copies/g)** | **Estimated serum viral load (copies/mL)** |
| --- | --- | --- | --- | --- |
| FEb1 | + | 34.36 | 898 | 222,512 |
| FGw | + | 35.59 | 377 | 120,558 |
| FDv | + | >40 | - | 9,399 |
| FBy | + | >40 | - | 31,922 |
| FOt | + | >40 | - | 30,086 |
| FAa1 | - | >40 | - | 0 |
| FBw | - | >40 | - | 0 |
| FGa1 | - | >40 | - | 0 |
| FHy | - | >40 | - | 0 |
| FHz | - | >40 | - | 0 |

## Discussion

African nonhuman primates can carry many types of RNA viruses, and are natural hosts for over 40 strains of SIV. Cross-species transmission of SIV from nonhuman primates to humans lead to the HIV/AIDS pandemic (Apetrei et al., 2004). HIV and SIV viral load are also the best predictor of AIDS progression in humans and macaques. As such, it is vital to characterize both the presence of SIV and the dynamics of SIV viral load in wild primate populations to better understand infection and transmission risks (Mellors et al., 1996; Staprans et al., 1999). However, preservation and detection of RNA viruses in noninvasively collected primate fecal samples can be challenging In this study, we tested multiple “field friendly” storage buffers and determined that DNA/RNA Shield was most effective in preserving SIV viral RNA in fecal samples. We then validated this approach in SIV-positive and negative sooty mangabeys from Emory NPRC.

SIV viral RNA was observed at the greatest abundances in all samples stored in DNA/RNA Shield at all times points and SIV concentrations as compared to all other buffers (with a single exception: RNA*later*, Week 12, Low SIV concentration) (**Figure 2c**). In two DNA/RNA Shield samples, we observed a viral RNA yield above expected (>100%) based on the amount that was spiked into each sample: Week 1, Medium SIV concentration – 155% yield; Week 1, SIV Low concentration – 105% yield. This may have been due to a non-homogenous distribution of SIV virions in these two sample aliquots.

RNA*later* has become one of the most widely used buffers for storing non-human primate fecal samples collected in the field; however, our results indicate that DNA/RNA Shield is a more effective choice for preserving viral RNA in fecal samples (**Table 3**). DNA/RNA Shield is advertised as being stable at room temperature for up to one month. At week 4, DNA/RNA Shield preserved between 76% and 86% of the SIV virions spiked into each sample at all concentrations. Notably, we observed a non-significant decrease in virion concentrations over time in DNA/RNA Shield at medium and low SIV concentrations. However, these samples were kept at room temperature for the duration of the study (12 weeks), which was well past the manufacturer’s recommended shelf stable timeframe (1 month). Had we followed manufacturer protocols, it is possible that virion concentrations would have stabilized to an even greater degree, rather than continuing to decline over time.

**Table 3:** Comparison of buffers for preservation of RNA viruses in fecal samples. ^*^Available from multiple manufacturers.

| Buffer | Manufacturer | Cost per mL | Fecal sample-to-buffer ratio (mg:mL) | Time shelf stable (at 25°C) | Percent yield (High concentration, Week 4) | Considerations | Examples of use in previous studies |
| --- | --- | --- | --- | --- | --- | --- | --- |
| **DNA/RNA Shield** | Zymo | ~\$1.90 | 100:1 | >30 days | 85.7% | Incompatible with some non-Zymo kits and workflows. | (Barrera-Avalos et al., 2022) |
| **RNAlater** | Thermo Fisher | ~\$1.10 | 200:1 | 1 week | 50.0% | Usable with most commercially available kits and workflows. | (Gopalakrishnan et al., 2021; Wille, Yin, Lundkvist, Xu, & Muradrasoli, 2018) |
| **Viral Transport Medium** | * | ~\$0.30 | 1000:1 | Not shelf stable | 20.9% | Samples must be frozen. | (Barrera-Avalos et al., 2022; Latorre-Margalef et al., 2014) |
| **95% Ethanol** | * | <\$0.10 | 1000:1 | Indefinitely | 0.1% | Shelf stable for DNA, but not shelf stable for RNA. | (Krafft et al., 2005) |

We also observed that a few Week 1 samples had moderately (but non-significantly) less SIV viral RNA as compared to subsequent timepoints (Weeks 4, 8, or 12). While the SIV virions should have experienced the least degradation at Week 1, and SIV viral RNA yield should have been the highest at this timepoint, it is possible that the extracted viral RNA in these samples degraded over time in the freezer. Week 1 viral RNA was stored frozen longer than samples from all other timepoints. These non-significant differences could also be attributed to minor variation in sample aliquoting and measurement.

Interestingly, both DNA/RNA Shield and RNAlater produced higher percentage yields of SIV viral RNA as compared to VTM, which was used as our “gold standard” (**Figure 2**). The CDC has recommended VTM for the storage of RNA viruses, such as SARS-CoV-2, for “efficient diagnosis” (Center for Disease Control and Prevention, 2021). However, similar to our results, previous studies comparing DNA/RNA Shield to VTM in SARS-CoV-2 testing have observed lower viral abundances (higher Ct values) in samples stored in VTM than those stored in DNA/RNA Shield. This indicates that DNA/RNA shield may be an even more effective alternative to the current gold standard for RNA virus preservation in biological samples (Barrera-Avalos et al., 2022).

We also observed consistently low viral RNA yield in samples stored in 95% Ethanol (less than 31% SIV viral RNA yield at all concentrations and time points). Ethanol is commonly used in non-human primate field studies for the long term storage of fecal samples, and it is therefore important to understand the limitations of this buffer. However, despite its limitations, we were able to successfully detect SIV in ethanol preserved samples at weeks 1, 4, and 8 (**Figure 2c**), indicating that RT-PCR is highly sensitive to the presence of SIV in ethanol, even at low SIV concentrations.

Based on the performance of DNA/RNA Shield in our laboratory experiment, we sought to validate these results using fecal samples from naturally infected SIV-positive and SIV-negative non-human primates. We did not detect SIV RNA in any of the SIV-negative mangabeys; however, we successfully detected SIV and quantified viral load in fecal samples from two of the five SIV-positive sooty mangabeys. These fecal samples both had low viral loads – less than 1000 copies SIV/g feces, which is considerably lower than viral loads typically found in serum (Ling et al., 2003; Palesch et al., 2018). Additionally, both positive fecal samples were associated with serum viral loads greater than 120,000 copies/mL. All negative fecal samples from SIV-positive mangabeys were associated with serum viral loads below 32,000 copies/mL. Sooty mangabey serum viral loads may range from less than 500 copies/mL to greater than 2×10^6^ copies/mL (Ling et al., 2003). This indicates that fecal viral shedding is lower in individuals with lower serum viral loads, making it more difficult to detect SIV in these individuals using noninvasive methods. The primary site of SIV/HIV viral replication is lymphoid tissue, where the virus enters and replicates within CD4+ T cells. These cells circulate throughout the body via the blood stream, making serum testing the most effective method for SIV/HIV testing. The gut also features an extensive array of lymphoid tissue known as gut-associated lymphoid tissue (GALT) in which SIV/HIV viral replication occurs, which can result in viral shedding into the feces (Veazey et al., 1998). Increased fecal viral shedding during HIV infection has been associated with gut dysfunction (Yolken et al., 1991). However, as natural hosts of SIV, sooty mangabeys do not progress to AIDS, and do not experience gut dysbiosis or inflammation during chronic SIV infection. They are therefore less likely to shed large amounts virus into the feces.

In conclusion, our results indicate that DNA/RNA Shield effectively preserved SIV viral RNA in fecal samples stored at room temperature for up to 3 months. However, given the decreases in SIV viral RNA yield that we observed (even between Weeks 1 and 4, which is within manufacturer guidelines for shelf-stable storage), samples should be shifted to a -80°C freezer as quickly as possible. These results also demonstrate that RT-PCR can be used to quantify SIV viral concentration in fecal samples, rather than simply report presence or absence of virus, which may help improve our knowledge of viral load and viral shedding in wild primates. Moreover, results from our validation cohort confirmed that DNA/RNA Shield preserved SIV viral RNA in fecal samples of naturally-infected non-human primate hosts, indicating the efficacy of this buffer for preserving RNA virus in feces.

## Materials and Methods

### Sample collection and preservation

A fresh fecal sample was collected from a single healthy SIV-negative mantled guereza colobus (*Colobus guereza)*, that was housed at the Columbus Zoo and Aquarium. The sample was immediately transported on ice to The Ohio State University College of Veterinary Medicine, homogenized, and aliquoted into multiple tubes with four different storage buffers: RNA*later* (Thermo Fisher), Viral Transport Medium (Medium 199 with 0.5% FBS), DNA/RNA Shield (Zymo Research), and 95% Ethanol (**Figure 1**). Fecal samples in each buffer were then inoculated with three concentrations of SIV virus: 300,000 virions/mL (high), 30,000 virions/mL (medium), and 3,000 virions/mL (low). These concentrations fall within the range of SIV concentrations detected in the plasma of naturally-infected SIV-positive primates (<500 copies/mL to greater than 2×10^6^ copies/mL (Ling et al., 2003)).

All samples were then stored at room temperature (25°C) for up to 12 weeks, with the exception of those in viral transport media (VTM) which were immediately stored at -80°C and considered our “gold standard” for sample preservation of RNA viruses (Abdallah et al., 2020). For many of these storage buffers, 12 weeks at room temperature is not recommended for optimal results, but we chose to extend our sampling to 12 weeks to mimic a long-term field study. Viral RNA was then extracted at four different time points (Week 1, 4, 8, and 12) using the QIAGEN RNeasy PowerMicrobiome Kit (Hilden, Germany). Negative controls were included for each buffer at each time point. Following extraction, viral RNA was frozen at -80°C.

### SIV plasmid linearization/standard curve

Plasmid DNA containing one copy of the SIV gene was linearized using EcoR1 (New England Biolabs, Ipswich, MA) following their restriction digest protocol (New England BioLabs, 2023). In brief, 2 μg of plasmid DNA was combined with 40 units of EcoR1, NEB buffer, and deionized water to a final volume of 50 μL. The resulting solution was incubated for one hour at 37°C. Following incubation, EcoR1 was then heat inactivated for 20 minutes at 65°C. The resulting linear DNA was then cleaned and concentrated with a commercially available kit (Zymo, Irvine, CA). The final DNA concentration was quantified via Qubit fluorometer (ThermoFisher, Waltham, MA) and used without further purification.

Following linearization, a standard curve was constructed via RT-PCR (Agilent, Santa Clara, CA) with copy numbers of the SIV gene ranging from 4.25 x 10^9^ copies to 85 copies using serial dilutions. RT-PCR was performed utilizing the S-GAG set of primers and probe (Cline et al., 2005)(Sigma-Aldrich, Burlington, MA) with a 1-step Master mix (ThermoFisher, Waltham, MA). The cycling conditions were 48°C for 10 minutes followed by 95°C for 10 minutes and 45 cycles of 95°C for 15 seconds and 60°C for 1 minute. The standard curve generated the following equation in which y = Ct value and x= viral copies: y = -1.42ln(x) + 39.443, R^2^ = 0.9978 (**Figure S1)**.

### Quantification of SIV viral load via RT-PCR

SIV viral RNA was amplified via RT-PCR using GAG primers and probe. Samples were run in triplicate on a 7500 Fast PCR instrument. RT-PCR cycling conditions were as follows: 10 minutes at 48°C; 10 minutes at 95°C; 15 seconds at 95°C and 1 minute at 60°C, for 45 cycles.

Negative controls for each buffer at each time point, and for the RT-PCR amplification process were included in the RT-PCR run. Triplicate cycle threshold (Ct) values for each sample were then averaged to estimate SIV viral load. Replicates with Ct values that exhibited greater than 3% variation were excluded from analysis, and all samples required at least two replicates within 3% variation to be included in analysis. The lowest Ct value associated with any negative control was 36.4; therefore, any true (non-control) sample that had a Ct value above 36.4 was excluded from analysis (**Table S1**). SIV virion concentrations and percent yield expected were then calculated based on Ct values and the standard curve described above. We then compared SIV virion concentrations between buffers and over time using Kruskal-Wallis one way analysis of variance followed by pairwise Dunn’s tests.

### Validation cohort

We collected blood and fecal samples from five SIV-positive and five SIV-negative sooty mangabeys (*Cercocebus atys*) housed at Emory National Primate Research Center (NPRC) in Atlanta, GA. Fresh fecal samples were collected opportunistically within 1 hour of defecation after individuals were isolated in cages for their annual veterinary exams. All fecal samples were placed in DNA/RNA Shield and stored at -80°C until RNA extraction. Concurrent blood samples were collected from the mangabeys during their annual exams to confirm SIV status. Viral RNA was extracted from fecal and blood samples using Zymo Quick-RNA Viral Kit. RT-PCR was run on all samples using the methods described above, and Ct values were used to quantify SIV viral load.

## Acknowledgements

This work was supported by an Ohio State University Infectious Diseases Institute Trainee Transformative Award (TC, RKS), an Ohio State President’s Research Excellence Accelerator award (VH, YV, WSM, TC). Tessa Cannon is also supported by an Ohio State University Presidential Fellowship. Samples collected from Emory National Primate Research Center were supported by ORIP/OD P51OD011132. The funders had no role in study design, data collection and interpretation, or the decision to submit the work for publication. We gratefully acknowledge Dr. Randy Junge and keepers at the Columbus Zoo and Aquarium for obtaining the fresh *Colobus guereza* fecal samples used in this study.

## Author contributions

Project conceptualization: TW, RKS, CM, YV, AS, WSM, VH

Virus preparation: RKS, AS

RNA extraction, RT-PCR: TW

Data processing and analysis: TW, CM

Interpretation and conclusions: TW, CM, VH

Manuscript writing: TW, VH

Manuscript editing: TW, RKS, CM, YV, AS, WSM, VH

